# A potent SARS-CoV-2 neutralizing human monoclonal antibody that reduces viral burden and disease severity in Syrian hamsters

**DOI:** 10.1101/2020.09.25.313601

**Authors:** Anna C. Fagre, John Manhard, Rachel Adams, Miles Eckley, Shijun Zhan, Juliette Lewis, Savannah M. Rocha, Catherine Woods, Karina Kuo, Wuxiang Liao, Lin Li, Adam Corper, Dilip Challa, Emily Mount, Christine Tumanut, Ronald B. Tjalkens, Tawfik Aboelleil, Xiaomin Fan, Tony Schountz

**Affiliations:** Department of Microbiology, Immunology, & Pathology, College of Veterinary Medicine and Biomedical Sciences, Colorado State University, Fort Collins, Colorado, USA 80523; AvantGen, Inc. San Diego, California, USA 92121; Department of Environmental & Radiological Health Sciences, College of Veterinary Medicine and Biomedical Sciences, Colorado State University, Fort Collins, Colorado, USA 80523

## Abstract

The emergence of COVID-19 has led to a pandemic that has caused millions of cases of disease, variable morbidity and hundreds of thousands of deaths. Currently, only remdesivir and dexamethasone have demonstrated limited efficacy, only slightly reducing disease burden, thus novel approaches for clinical management of COVID-19 are needed. We identified a panel of human monoclonal antibody clones from a yeast display library with specificity to the SARS-CoV-2 spike protein receptor binding domain that neutralized the virus *in vitro*. Administration of the lead antibody clone to Syrian hamsters challenged with SARS-CoV-2 significantly reduced viral load and histopathology score in the lungs. Moreover, the antibody interrupted monocyte infiltration into the lungs, which may have contributed to the reduction of disease severity by limiting immunopathological exacerbation. The use of this antibody could provide an important therapy for treatment of COVID-19 patients.

## Introduction

The emergence of a novel coronavirus disease (COVID-19), caused by severe acute respiratory syndrome coronavirus 2 (SARS-CoV-2), was first described in December 2019 in Wuhan, China^1^. Symptoms include cough, dyspnea, chest tightness and fever, plus occasional adverse gastrointestinal (GI) disturbances. In a vulnerable subset of patients, the disease often progresses to an atypical pneumonia with high morbidity and mortality rates^2–6^. Mortality based on case fatality rates have ranged from early estimates of 3.9% in Hubei province, China, to 0.9% in populations from countries with widespread testing. Recent estimates of infection fatality rates (IFRs) are between 0.6% to 1% and overall indicate that COVID-19 has a 10-fold greater mortality than seasonal influenza^6–10^. Alarmingly, the IFR is considerably higher (5.6%) in individuals ≥65 years of age where the prognosis for those hospitalized patients ranges from guarded to poor; they frequently require long periods of breathing support, with mortality rates of 20-30%^6,11^. Consequently, the effect on healthcare resources is of major concern and there is an increasingly unmet medical need for effective therapies, particularly for older patients with comorbidities, including atherosclerosis, hypertension and diabetes mellitus, to prevent lethal pneumonia.

Cellular infection by coronaviruses is mediated by the viral homotrimeric spike glycoprotein binding to specific host cell receptors. Each protomer consists of an S1 domain, which mediates receptor binding, and an S2 domain that mediates membrane fusion and cell infection following conformational changes induced by host cell receptor binding ^12–14^. The receptor for SARS-CoV, SARS-CoV-2 and HCoV-NL63 is angiotensin converting enzyme 2 (ACE2), which is expressed on mucosal epithelia of the upper respiratory tract, bronchioles, lungs and the GI tract^15–19^. Convalescent serum and purified antibodies from recovering patients have yielded promising results in past viral outbreaks^20–26^, and this approach may also be promising for SARS-CoV-2.

One key challenge in the development of these antibodies is how to prevent viral escape whereby variant mutations nullify antibody binding to the RBD epitope. Multiple variants of SARS-CoV-2 have emerged, some of which have mutations in the RBD of the S protein and are predicted to have increased binding to ACE2^27–30^. The best neutralizing antibodies are likely to recognize epitopes with high sequence conservation due to structural or functional constraints. The RBM binding site to ACE2 is such an example because sequence divergence within this region is constrained to the prerequisite that binding to ACE2 must be maintained.

An analysis of the crystal structures of the ACE2 - RBD SARS-CoV complex^31^ and the ACE2-SARS-CoV-2 RBD complex^29,32^ confirms that eight of the key RBM contact residues are conserved between these respiratory coronaviruses, although the SARS-CoV-2 RBM forms a more compact interface with ACE2 than that formed by SARS-CoV RBD and has a higher affinity for ACE2^29,32^. Many of the previously identified neutralizing antibody clones against SARS-CoV were shown to block S1 binding to its ACE2 receptor by binding to epitopes located within the RBM^21–24,33,34^. However, not all monoclonal antibodies isolated previously against the SARS-CoV RBD recognize the SARS-CoV-2 spike protein in the context of the full-length spike homotrimer^35,36^. Similarly, recent studies evaluating anti-SARS-CoV-2 antibodies isolated from convalescent patients indicate only a minor subset of the clones have the requisite viral neutralizing activity^35,36,46^.

Together, these studies indicate that there is an urgent need for SARS-CoV-2-specific human neutralizing antibody therapeutics that have proven ability to both neutralize SARS-CoV-2 viral host cell infection *in vitro* as well as the appropriate activity *in vivo*. The Syrian hamster has been shown to be a suitable model for SARS-CoV-2 infection given the similarity of clinical signs and lung pathology to human disease in the first few days of infection^37–39^. However, to date, there has been only a gross histological analysis of the lung pathological changes following infection and the impact of SARS-CoV-2 neutralizing antibody clones on lung immune infiltrates has yet to be fully assessed. In this study, we describe an anti-SARS-CoV-2 spike RBD clone isolated from a rationally designed, fully human antibody library that bound to native spike protein. This potent antibody could block the interaction of the spike protein with ACE2 and could also block SARS-CoV-2 infection of Vero E6 cells and the resultant cytopathic effect. This clone was then tested in a pilot therapeutic study in Syrian golden hamsters (*Mesocricetus auratus*) and was shown to reduce viral load and ameliorated the severity of bronchointerstitial pneumonia. Detailed histological and image analysis suggests that macrophages play a key role in the lung response to SARS-CoV-2 infection in hamsters.

## Materials and Methods

### Production and purification of antibody clones

The DNA inserts encoding the light and heavy chains of clone AvGn-B and its variants were separately cloned into different expression vectors carrying the constant regions of human IgG1 heavy chain and the kappa chain. The heavy and light chain plasmids were co-transfected into Expi293 suspension cells. After 5-7 days, secreted antibody was then purified from the culture supernatants by protein A chromatography, buffer-exchanged into PBS pH 7.4 and its concentration determined by Nanodrop A280 assay. The quality of the IgG was assessed by SDS-PAGE and by HPLC. Endotoxin levels were tested using a Limulus amoebocyte lysate (LAL) assay (ThermoFisher Scientific, cat # A39552).

### Determining affinity and receptor binding inhibitory activity of clone AvGn-B and its variants

The wells of Immulon 2 HB ELISA 96-well plates were coated with AvGn-B or its variant IgGs at 4° C overnight. The wells were blocked, washed, then serial dilutions of the monomeric biotinylated RBD-antigen added to the wells and incubated for 1 h at room temperature. Bound antigen was then detected with HRP-labeled streptavidin. The OD readings were plotted against concentration using Prism software (Graphpad CA) for curve fitting and determination of the apparent K_D_ value.

The IC_50_ values for inhibition of RBD binding to the ACE2 receptor were determined by pre-mixing a serial dilution of antibody with 2 nM of biotinylated SARS-COV-2-RBD and incubating for 30 min before adding to wells coated with recombinant ACE2-Fc fusion protein (residue Gln18-Ser760 fused to human Fc [Kactus BioSystems, Cat# ACE-HM401]). After 1 h, bound RBD was detected using HRP-labeled streptavidin and the OD readings plotted against antibody concentration using Prism software.

Real time kinetics were measured by bilayer interferometry (BLI) at a temperature of 30° C using a Gator™ system (Probe Life Inc, CA). After establishing a baseline with filtered Q-buffer (PBS with 0.2% BSA and 0.02% Tween-20), IgG1 at 1 μg/mL (200μL per well) was captured onto six ProtG probes until a wavelength shift of approximately 1 nm was observed on the Gator sensorgram. After reestablishing baseline, binding of biotinylated SARS-CoV-2 RBD (Kactus Biosystems; Cat# COV-VM4BDB) analyte (200 μL per well) to captured antibody was measured via separate association [six concentrations (1 per well) consisting of a 3-fold serial dilution range of 138.9 - 0.57 nM] and dissociation steps. Non-specific binding events, to the probe or to an irrelevant control antibody, were monitored in two separate wells and the reference subtracted from each test antibody sensorgram. Antibody K_on_, K_off_ and K_D_ values were determined using Gator software through global fitting of the data to a 1:1 binding model.

### Assessing antibody binding to native spike protein

A vector encoding a full-length spike protein sequence encompassing residues 13-1273 of UniProtKB accession number P0DTC2, that was fused to green fluorescent protein (GFP) as a reporter (Sino Biologicals, Cat# VG40590-ACG) was used to transiently transfect 293F cells using the FreeStyle™ 293F Expression System (ThermoFisher Scientific, cat# K90001). After 2 days, when a robust GFP signal was observed, aliquots of the transfected cells and untransfected 293F cells were washed in ice cold phosphate-buffered saline containing EDTA and 0.5% BSA, pH 7.4 (PBSM) then incubated in the presence of AvGn-B, isotype control antibody or ACE2-Fc for 45 mins with rotation at 4° C, washed 3 times in ice-cold PBSM and then bound antibody detected with goat anti-human kappa-A647 (SouthernBiotech, cat# 2060-31) After washing, cell-bound Alexa-657 was analyzed using an IntelliCyt^®^ iQue Screener PLUS and ForeCyt Software (Sartorius).

### Determining the viral neutralization activity of antibody clones in vitro

Vero E6 cells were plated overnight in 96-well plates at 20,000 cells per well. Antibodies were diluted in Complete DMEM and serially diluted 1:3 resulting in a 12-point dose response dilution series run in 4 or 8 replicates. The dilutions of antibody were incubated with 100 TCID_50_ per 50 μL of SARS-CoV-2 (strain 2019-nCoV/USA-WA1/2020) for 1 h and added to the assay plates. The plates were incubated for 3 days at 37° C, 5% CO_2_ and 95% relative humidity and the inhibitory effects of the antibodies were assessed.

### Study design for animal infections

Approval of the study protocol was obtained from the Colorado State University Institutional Animal Care and Use Committee (protocol 993). Male Syrian hamsters (n=20, 10 weeks of age, obtained from Charles River Laboratory). The animals were held in the CSU animal facility and provided access to standard pelleted feed and water *ad libitum* prior to being moved into the biosafety level 3 facility for experimental challenge. Of the 20 hamsters, 18 were intranasally infected with 2.5×10^4^ TCID_50_/mL equivalents of SARS-CoV-2 (strain 2019-nCoV/USA-WA1/2020) and divided into treatment groups as follows: “AvGn-B High” (2.5 mg AvGn-B) (n=5), “AvGn-B Low” (1 mg AvGn-B) (n=5), “Untreated” (no antibody) (n=6), and “Ab Control” (2.5 mg isotype control IgG) (n=2). Two uninfected hamsters received 2.5 mg AvGn-B (termed “Uninfected”). Each animal was dosed intraperitoneally with corresponding treatment at 24 and 72 hours post dosing (hpi). At 24, 48, 72, 96, and 120 hpi, each hamster was weighed and assessed for presence of clinical signs (lethargy, ruffled fur, hunched back posture, nasolacrimal discharge, and rapid breathing). At 120 hpi (5 dpi), each hamster was anesthetized with isoflurane and then euthanized via cardiac exsanguination and blood was collected.

Weight loss (calculated as percentage decrease from 0 dpi weight) and viral RNA load in lung were compared between treatment groups in Prism using multiple t-tests and Mann-Whitney U tests, respectively. *p*<0.05 was considered significant.

### RNA extraction and qRT-PCR

Swabs in viral transport medium were vortexed thoroughly and centrifuged to pellet cellular debris. RNA was extracted from swab supernatant using the QiaAmp Viral RNA Mini Kit (Qiagen, Cat #1020953) according to the manufacturer’s instructions. Lung tissue was homogenized with a Qiagen TissueLyser LT and RNA extracted from supernatant using Qiagen RNeasy Mini Kit (Qiagen, Cat #74104) following manufacturer instructions. The Realtime Ready RNA Virus Master (Roche, Cat # 05619416001) was used to amplify viral RNA with the following primers/probe: Forward: 5’-ACAGGTACGTTAATAGTTAATAGCGT-3’, Reverse: 5’-ATATTGCAGCAGTACGCACACA-3’, Probe: 5’-FAM-ACACTAGCCATCCTTACTGCGCTTCG-BBQ-3’ as previously described^40,41^. A 2019-nCoV_E positive control plasmid (Integrated DNA Technologies, Cat #100006896) was used to generate a standard curve for copy number quantification.

### Histopathology and immunohistochemistry

Lungs, tracheobronchial and hilar lymph nodes, thymus, esophagus, heart and liver from 20 hamsters labeled with ear tags 26-45 were extirpated *en bloc* and fixed whole in 10% neutral buffered formalin for at least 3 days to ensure virus inactivation prior to transfer to the CSU Veterinary Diagnostic Laboratory for trimming.

Four transverse whole-lung sections were stained with H&E or processed for IHC. Sections, 5 μm thick, were subjected to heat-induced epitope retrieval performed online on a Leica Bond-III IHC automated stainer using bond-epitope retrieval solution. Antibodies to SARS-CoV-2 nucleocapsid protein (mouse, 1:500), pancytokeratin, factor-VIII and ionized calcium binding adaptor molecule (IBA-1) (Leica Biosystems) or negative control slides primary antibody was replaced by a rabbit non-specific IgG isotype negative control antibody for 20 minutes. Labeling was performed on an automated staining platform. Fast red was used a chromogen and slides were counterstained with hematoxylin. Immunoreactions were visualized and blindly scored by a single pathologist. In all treated categories, reactive lung sections incubated with primary antibodies were used as positive immunohistochemical control. Negative control sections were incubated in diluent composed of Tris-buffered saline with a carrier protein and homologous nonimmune serum. All sequential steps of the immunostaining procedure were performed on negative controls following incubation.

### Immunofluorescence Staining and Imaging

Paraffin embedded tissue sections were stained for SARS-CoV-2 nucleocapsid protein (1:500) and ionized calcium binding adaptor molecule 1 (Abcam; ab5076; 1:50) using a Leica Bond RX^m^ automated staining instrument following permeabilization using 0.01% Triton X diluted in Trisbuffered saline (TBS). Blocking was performed with 1% donkey serum diluted in TBS. Sections were stained with DAPI (Sigma) and mounted on glass coverslips in ProLong Gold Antifade mounting medium and stored at ambient temperature until imaging. Images were captured using an Olympus BX63 fluorescence microscope equipped with a motorized stage and Hamamatsu ORCA-flash 4.0 LT CCD camera. Images were collected with Olympus cellSens software (v 1.18) using an Olympus X-line apochromat 10× (0.40 N.A.), 20× (0.8 N.A.) or 40× (0.95 N.A.) air objectives, or Uplan Fluor ×100 oil immersion (1.3 N.A.) objective. Regions of interest (ROIs) were drawn around the perimeter of each tissue section and intensity thresholds were kept constant across groups analyses were performed blinded. Co-localization and intensity measurements were obtained using the Count and Measure feature of cellSens.

### Digital Image Analysis

All lung sections were digitalized through 4×, 10×, 20× and 40× objectives on a Nikon Eclipse 80i microscope and Nikon Digital Sight DS-Fl1 camera (Nikon instruments, USA). Quantitative and qualitative microscopic analyses were determined using micrographs and morphological methods to quantify pulmonary lesions with the use of NIS-Elements (Nikon, Americas, USA). Stained slides with H&E were manually screened for pattern recognition of affected versus unaffected pulmonary parenchyma. At least 6 representative ROIs were defined (1088 x 816 μm) and subjectively delineated (annotated) by a single trained veterinary pathologist based upon hypercellularity and consolidation of alveolar spaces. Manual annotation was compiled for digital quantification of affected areas of the lung to compare individual hamsters. Data were expressed as mean (±SEM). Statistical analyses were performed using one-way ANOVA multiple comparison test. *p* values less than 0.05 were considered significant. Horizontal whole-lung slides of consecutive planes of SARS-CoV-2-infected and antibody-treated lungs were digitalized by Olympus histoslide scanner to create image data files that can be used for further digital image analysis (DIA). Establishment of the algorithm started with the definition of lesional and non-lesional components (classes) according to Table 1.

**Table 1.** Scoring classification of hamster lung pathology.

| **Class** | **Designation** | **Characteristic alterations** |
| --- | --- | --- |
| **1** | Affected lung area<br>“blue” and “red” for<br>cellular density | Intra-bronchiolar, intra-alveolar, peribronchiolar<br>and perivascular inflammation or alveolar septal<br>thickening, edema and hemorrhage |
| **2** | Unaffected lung<br>tissue “light blue” | Normal lung parenchyma |
| **3** | Background “grey” | Intravascular erythrocytes, blood vessel walls,<br>bronchiolar mucosa |
| **4** | Glass “white” | Glass |

### Quantification of total nuclei, inflammatory cells and consolidation

The precise digital quantification of the total affected pulmonary parenchyma as well as counting of inflammatory cells per area (ROI, 1mm^2^) in histological section was determined. A digital montage was compiled at 100× magnification to include the four tissue classes that were differentially characterized (Table 1) to establish algorithm classifier. Affected ROIs were subsequently automatically identified using Olympus cellSens software by quantifying whole-lung mounts scanned from each hamster for total number of nuclei or nucleated cells (to exclude erythrocytes) stained with hematoxylin. Sections labeled by immunofluorescence for IBA-1 and SARS-CoV-2 were analyzed using the Count and Measure Module of cellSens. The algorithm predominantly extracted multispectral information from images with additional DIA processing using spatial, logical and threshold separators after manual annotations. All four classes were accurately identified as expected by a board-certified pathologist who was blinded to experimental groups of hamsters.

## Results

### Isolation of an anti-SARS-CoV-2 RBD antibody clone that blocks receptor binding and can neutralize SARS-CoV-2 activity

A rationally designed fully human antibody library displayed on yeast was screened by magnetic bead sorting (MACS) followed by two rounds of fluorescence activated cell sorting (FACS) using recombinant SARS-CoV-2 RBD (Arg319-Asn532 with a His-tag and Avi-tag at the C-terminus; Kactus BioSystems, Cat# COV-VM4BDB) as antigen to enrich for yeast clones expressing antibody clones that bound the RBD antigen. The harvested cells were then subjected to a third round of FACS selection whereby the enriched yeast pool of RBD-binders was incubated in the presence of 50 nM biotinylated RBD and 100 nM human ACE2-human Fc fusion protein. Bound RBD was then detected with PE-labeled streptavidin and captured ACE2-human Fc was detected by Dylight 647 conjugated goat anti-human Fc. Clones that bound biotinylated RBD in a manner that prevented binding of ACE2-Fc to antigen were collected as those displaying potential inhibitory antibody clones that block the RDB-receptor interaction. After a final round of positive selection of clones that bound SARS-CoV-2-RBD, the harvested pools were plated for analysis of individual clones.

Individual antibody clones were tested for their abilities to block SARS-CoV-2-RBD binding to ACE2 using a competition ELISA and tested for their ability to bind to native SARS-CoV-2 spike protein expressed as a GFP fusion protein by transfected 293 cells compared to binding activity to non-transfected 293 cell controls. Those antibody clones that blocked the interaction of the RBD with ACE2 and bound to native spike protein were then tested for neutralization of SARS-CoV-2 in a cytopathic effect (CPE) assay with Vero E6 cells.

Clone AvGn-B was identified as the most potent in these assays. It exhibited an apparent affinity by ELISA for SARS-CoV-2 RBD of 0.17 nM (Fig. 1A), an apparent IC_50_ value for blocking RBD binding to ACE2-Fc of 2.2 nM (Fig. 1B), and an apparent K_D_ for binding to native spike protein expressed by 293 cells of 1.02 nM compared to 10.9 nM for ACE2-Fc (Fig. 1D). Real time binding kinetics of AvGn-B to the isolated RBD recombinant protein measured by BLI indicated a K_D_ of 0.37 nM (Fig. 1C). When tested for its ability to neutralize SARS-CoV-2 infectivity, Clone AvGn-B exhibited 100% protection from SARS-CoV-2 induced cell death down to 0.017 μg/mL. Using a colorimetric assay for quantitation of cell death, AvGn-B exhibited an IC_50_ value of that ranged from 0.008 μg/mL (experiment with 4 replicates) to 0.0054 μg/mL (experiment with 8 replicates) in the CPE assay (Fig. 1E).

**Figure 1.**
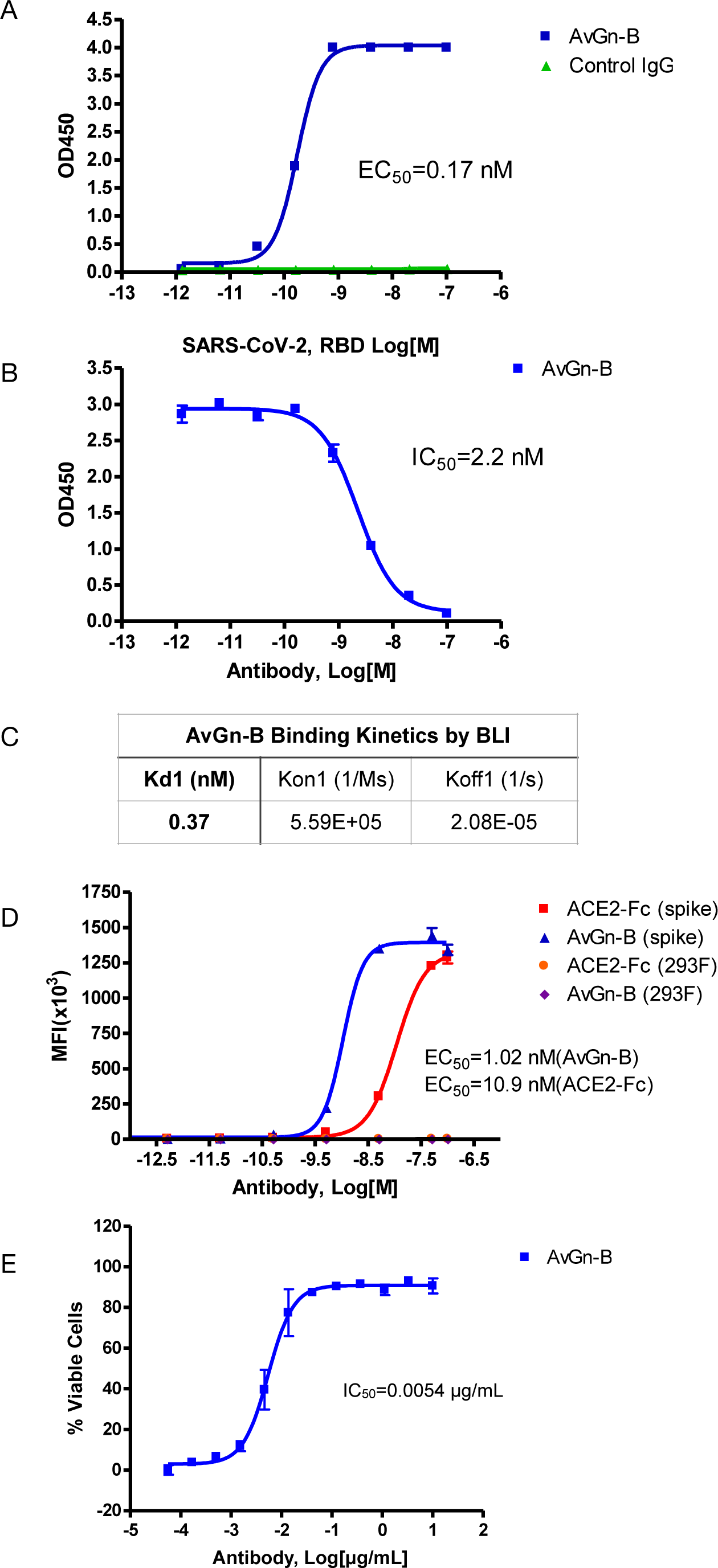
Properties of clone AvGn-B. **A.** Binding curve of biotinylated RBD to the virus neutralizing AvGn-B clone by ELISA. **B.** IC_50_ curve of the AvGn-B for competing binding of SARS-CoV-2 RBD to immobilized ACE2-Fc receptor. **C.** Binding kinetics of RBD to AvGn-B measured by BLI, using an Octet system. **D**. Binding curve of clone AvGn-B compared to ACE2-Fc to 293 cells expressing full-length SARS-CoV-2 spike protein as a GFP-fusion protein. **E.** EC_50_ curve for the ability of clone AvGn-B to neutralize viral-induced cell death in the CPE assay. Curve fitting in A, B, D and E was performed using Prism software.

### Evidence for SARS-CoV-2 neutralization *in vivo*

At 2 dpi, infected hamsters appeared quiet and began to progressively lose weight over the course of the study (up to ~13% reduction in untreated animals). There was no significant reduction in weight loss associated with AvGn-B treatment (*p*>0.05) (Fig. 2A). None of the hamsters in the study died or met euthanasia criteria prior to study termination at 5 dpi. There was a significant reduction in viral RNA in the lungs of hamsters treated with AvGn-B (both 2.5 mg and 1 mg doses) compared to those that were untreated (*p* = 0.0173 and 0.0303, respectively) (Fig. 2B).

**Figure 2.**
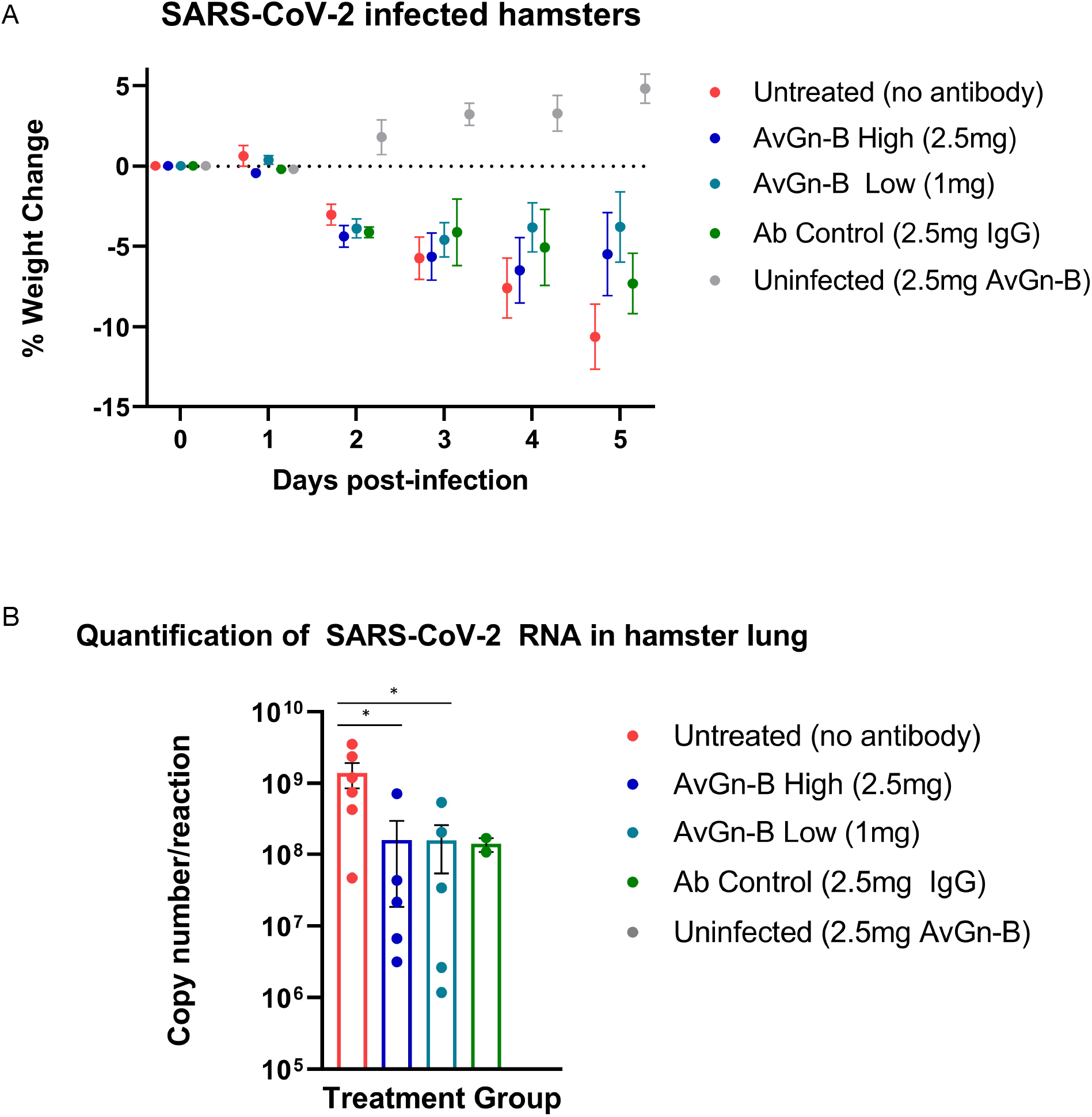
Clinical presentation and viral load in the lungs of Syrian hamsters dosed with AvGn-B following SARS-CoV-2 infection. **A.** Hamster body weights were recorded daily (0-5dpi), and weight loss was defined as percentage loss from 0dpi. Hamsters were separated by treatment group and weight loss was analyzed using multiple t-tests in Prism GraphPad for each day (p>0.05). **B.** Viral load (gene copy number/reaction) in infected hamster lung was compared between treatment groups (analyzed by Mann-Whitney U Test in Prism GraphPad) (\**p*<0.05).

### Pathology and immunohistochemistry results

Lungs from the 2 uninfected and the 2 Ab Control hamsters (treated with 2.5 mg control isotype IgG) did not show significant bronchiolar or parenchymal inflammation. Distal trachea and main stem bronchi contained microscopic hemorrhages and there were variable degrees of iatrogenic atelectasis, especially in cranial and middle portions of the lungs. Occasional peribronchiolar lymphoid follicles of moderate cellularity were prominent around bronchioles. Other organs, namely, tracheobronchial and hilar lymph nodes, esophagus, heart and kidneys were within normal histologic limits. In virus-infected, untreated hamsters (Untreated group), approximately 50-75% of lung parenchyma, especially the inner parenchyma (i.e., peribronchial lung tissue), was consolidated, dark red and heavier than outer inflated lung parenchyma. Lungs from negative control IgG1-treated hamsters (Ab Control group) did not show any consolidation or significant parenchyma inflammation (Fig. 3A). Massive cellular infiltrates of predominantly macrophages, 60-70%, intermixed with neutrophils, lymphocytes and plasma cells partially filled the lumina of terminal bronchioles and adjacent alveolar and perivascular spaces to varying degrees. In the most affected lobes, there was near complete obliteration of air spaces, which were manually annotated (Fig. 3B). Multifocally, bronchiolar epithelium and regenerative alveoli contained multinucleate giant syncytial cells that stained positive for pancytokeratin. Marked reduction in inflammatory cell infiltration into pulmonary parenchyma was observed in the AvGn-B High group where only 10-15% of total lung parenchyma was involved (Fig. 3 C and D). Total areas of lung parenchyma and ROI were annotated for cell counts (Fig. 3 E and F). Near the pleural surface and around primary bronchi, there were scattered alveoli lined by increased numbers of type II pneumocytes indicative of regeneration. Main pulmonary arterial vessels were lined by several rows of monocytes including scattered multinucleate giant cells (Fig. 4A and B). Endothelial cells lining affected vessels show multifocal loss due to sloughing of individual endothelial cells or vacuolation with crowding of macrophages along defective intima. In affected vessels, there was multifocal histiocytic infiltration of tunica media, which showed apoptosis of medial smooth muscle fibers (leukocytoclastic vasculitis) (Fig. 4C). Viral antigen was evident in cytoplasm of luminal monocytes and mural macrophages (Fig 4D). Factor VIII (endothelial marker showed disruption of continuity of the tunica intima (Fig. 4E) and IBA-1 staining proved the monocytic origin of the luminal multinucleate cells (Fig. 4E). Overall size of tracheobronchial and hilar lymph nodes increased 2-3-fold forming densely cellular lymphoid follicles. Adipose tissue at the base of the heart, surrounding great vessels showed multinodular infiltrates of moderate numbers of macrophages, lymphocytes and plasma cells with only a few neutrophils. Cellular infiltrates and intravascular monocytes moderately diminished in numbers in the AvGn-B Low group where only 25-35% of pulmonary parenchyma was involved. Residual inflammation was mainly observed in bronchial tree and large pulmonary arteries. No significant pathology or immunostaining was seen in the rest of parenchymatous organs.

**Figure 3.**
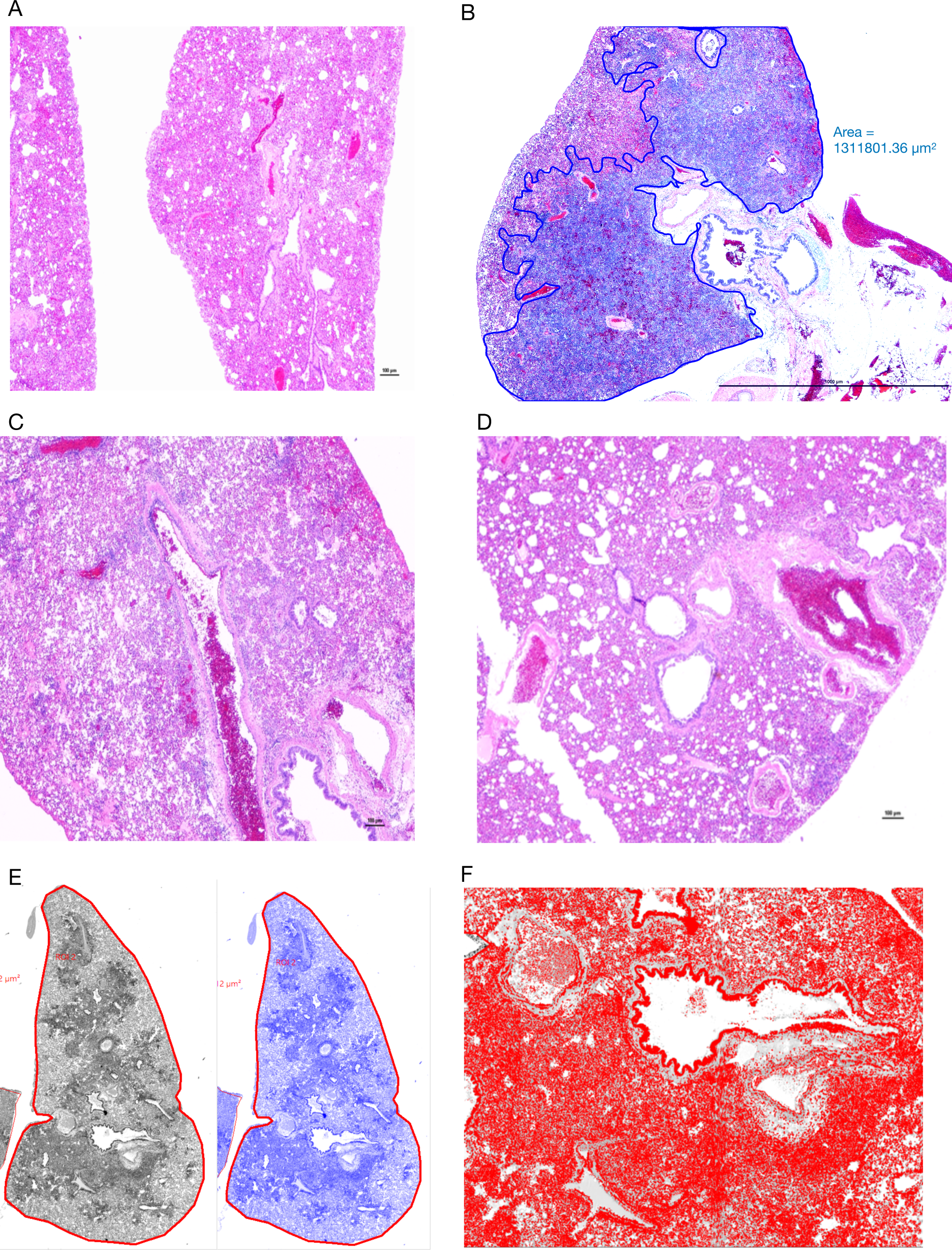
Manual annotation and comparison of histological scores of bronchointerstitial pneumonia in control and treated hamsters. **A.** Lungs from an uninfected hamster were visualized and scored for lack of significant hypercellularity. **B**. Affected parenchyma of infected hamsters (Untreated) characterized by intense hypercellularity is manually annotated (blue) excluding unaffected lung parenchyma, main stem bronchi and hilar adipose tissue. **C**. Lung section from a control antibody treated hamster (Ab Control) is showing marked reduction in total cellularity or inflammation. **D.** Lung section from a low-dose AvGn-B antibody-treated hamster (AvGn-B Low) showing very significant reduction of inflammation reducing the total area of affected pulmonary parenchyma. **E.** After manual annotation depicted in B and based on the generated montage including training set and threshold, the automated classifier to discriminate lesional from non-lesional parenchyma was generated in grey and then in blue. **F**. After manual annotation of affected lungs, all of the defined classes in preset ROI, *i.e*., inflammatory foci characterized by infiltration of inflammatory cells into alveolar spaces were identified to generate total number of nuclei (hematoxylin and DAPI-stained nuclei) that were accurately recorded.

**Figure 4.**
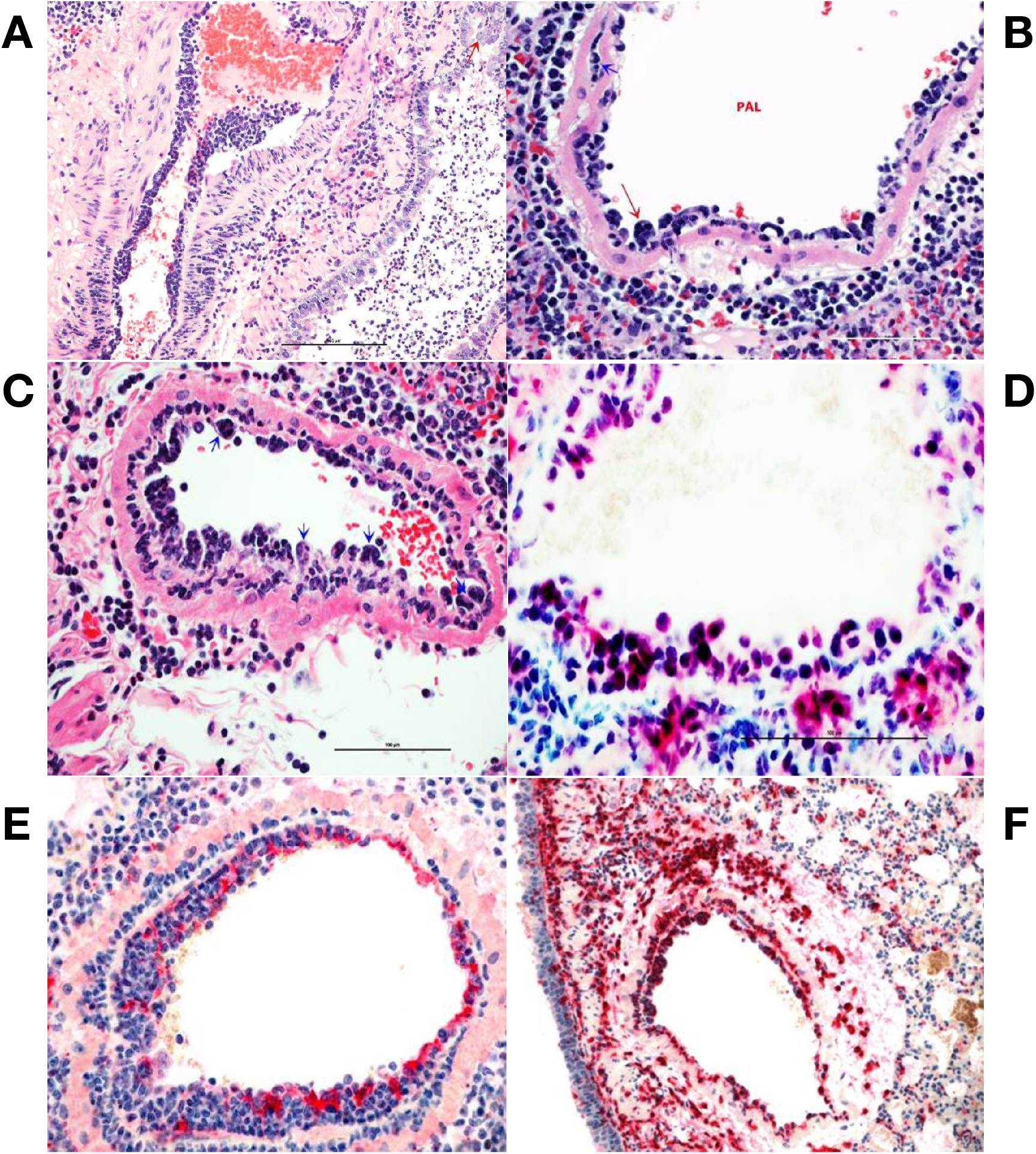
Monocytosis and histiocytic arteritis in SARS-CoV-2-infected and untreated hamsters. **A.** Major branches of pulmonary artery contain numerous monocytes adhered to endothelial cells. **B.** The luminal monocytes in pulmonary artery lumen (PAL) include multinucleate cells attached to the luminal surface of endothelium (red arrow) and the vessels is surrounded by many macrophages. **C.** Close-up of pulmonary artery tunica intima showing endothelial-adherent monocytes some of which are subintimal (blue arrows) or migrating through the wall of the vessels, leukocyoclastic vasculitis. **D.** Luminal monocytes and perivascular macrophages showing immunoreactivity against SARCoV-2 nucleocapsid. **E.** Staining the vessels intima with endothelial marker, factor VIII show discontinuity of the tunical intima by migrating monocytes. **F.** Most of the cells crowding tunica intima are showing strong immunoreactivity for the macrophage marker, CD204.

### Analysis of macrophage infiltration in lung tissue of animals exposed to SARS-CoV-2

Lungs were examined for the extent of macrophage infiltration and SARS-CoV-2 viral load at 5 days post infection by histopathological examination and by immunofluorescence imaging (Fig. 5). Whole mount sections of paraffin-embedded lung tissue were stained with hematoxylin and eosin and brightfield grayscale images were collected using a microscope equipped with a scanning motorized stage (Fig. 5A-E). Hematoxylin-positive macrophage soma were rendered as focal points within the regions of interest to calculate the percent hypercellularity of tissue following infection with SARS-CoV-2. By 5 dpi, lung tissue showed extensive infiltration of macrophages (Fig. 5A) that was decreased in dose-dependent fashion by treatment with AvGn-B (Fig. 5B, 5C). Treatment of uninfected hamsters with AvGn-B alone did not increase macrophage infiltration, whereas co-treatment with IgG control Ab during infection with SARS-CoV-2 (Ab Control group) still resulted in marked infiltration of macrophages (Fig. 5E). The percent of total lung area displaying macrophage hypercellularity was quantified in Fig. 5N, and reflected the observed decrease in total hypercellular lung area in animals infected with SARS-CoV-2 and treated with increasing concentrations of AvGn-B. These findings were confirmed by co-immunofluorescence imaging for macrophages (IBA-1^+^ cells) and SARS-CoV-2 (Fig. 5F-M). High-resolution montages scans of whole mount lungs revealed pan-lobular replication of SARS-CoV-2 and largely centrilobular infiltration of IBA-1^+^ macrophages. Treatment with AvGn-B at 2.5 mg resulted in a marked decrease in the presence of IBA-1^+^ macrophages throughout the lung. Quantification of IBA-1 intensity (Fig. 5O) and the number of IBA-1^+^ cells co-localizing with SARS-CoV-2 (Fig. 5P, representing viral phagocytosis) revealed similar trends and indicated that animals infected with SARS-CoV-2 and treated with 1 or 2.5 mg AvGn-B had marked decreased in infiltration of IBA-1^+^ macrophages and a corresponding decrease in lung tissue pathology.

**Figure 5.**
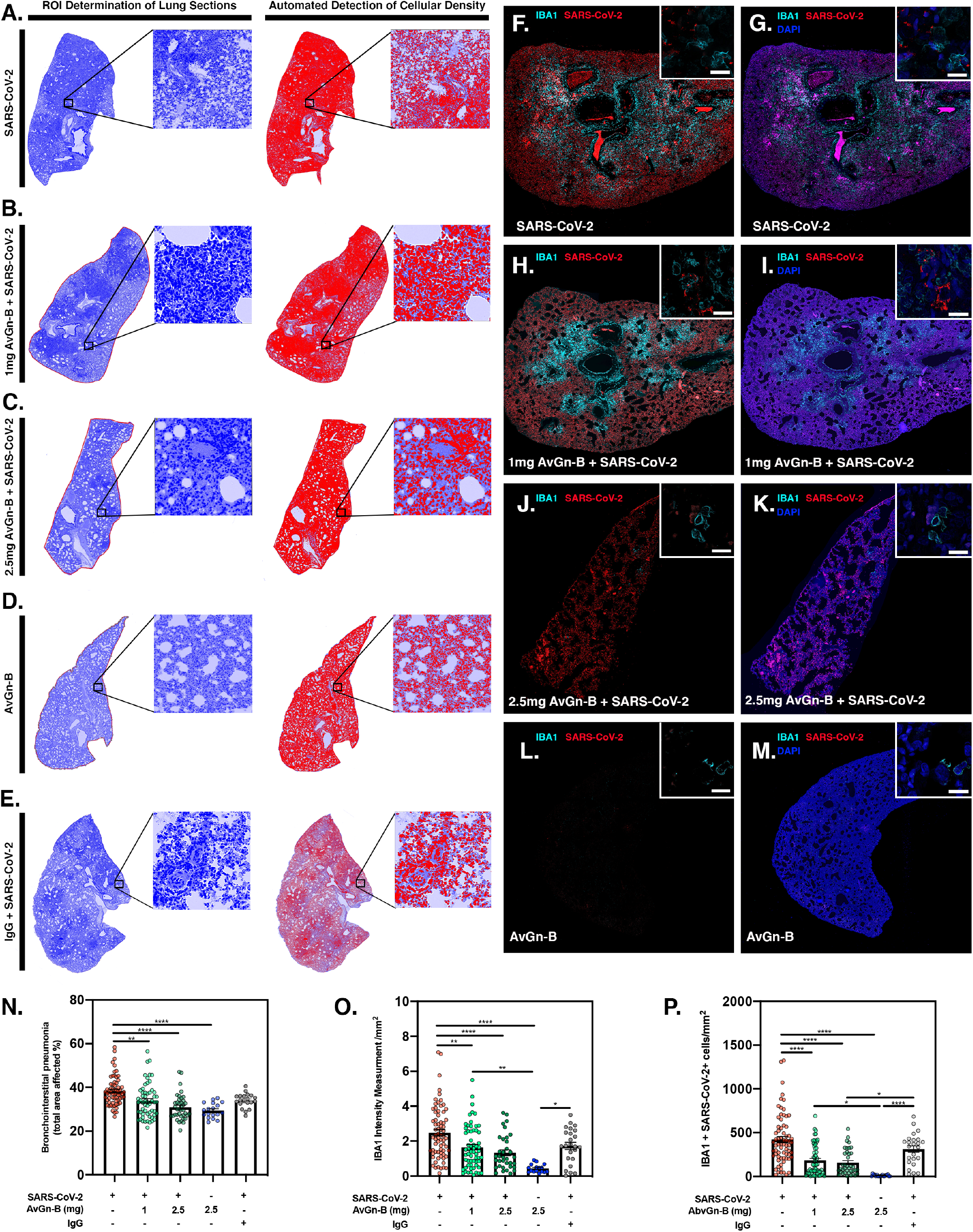
Reduction of SARS-CoV-2 and macrophage infiltration by AvGn-B treatment. **A-E.** Overall hypercellularity within lung tissue, resulting in pathological bronchointerstitial pneumonia, was determined using ROI delineations on hematoxylin and eosin stained sections for all groups. **N.** Quantification of the total area of the lung tissue affected with bronchointerstitial pneumonia was conducted using automated focal point determination within ROIs following manual thresholding. **O.** The total IBA-1+ monocyte lineage cellular intensity was quantified in the lung tissue (for each experimental group, as depicted in the high resolution (white bar in each high resolution image represents 100 μm) and 10× immunofluorescent montage images (**F-M**). **P.** Co-localization of SARS-CoV2 (red) and IBA-1 (cyan) was also quantified within the SARS-CoV2, SARS-CoV2+AvGn-B 1mg, SARS-CoV2+AvGn-B 2.5mg groups (\**p*<0.05, \*\**p*<0.01,\*\*\**p*<0.001, \*\*\*\**p*<0.0001; N=5 hamsters/group) as well as in the uninfected+AvGn-B 2.5 mg and SARS-CoV-2+IgG groups (\**p*<0.05, \*\**p*<0.01,\*\*\**p*<0.001, \*\*\*\**p*<0.0001; N=2 hamsters/group).

## Discussion

To date, few treatment options exist for COVID-19 patients. Identification of therapeutic monoclonal antibodies for COVID-19 would reduce its morbidity and mortality. Screening a set of rationally designed fully-human antibody libraries displayed on yeast led to the discovery of a handful of clones that exhibited neutralizing activity against SARS-CoV-2. The clone with highest *in vitro* affinity to SARS-CoV-2 RBD, AvGn-B, was also shown to reduce disease burden in SARS-CoV-2 infected Syrian hamsters, illustrating its potential as a therapeutic agent for use in COVID-19 patients.

AvGn-B exhibited potent activity, with an IC_50_ of 0.005-0.008 μg/mL, which compares favorably with other monoclonal antibodies shown to neutralize SARS-CoV-2^42–46^. While a few recent studies have shown the disease ameliorating effects of administering neutralizing SARS-CoV-2 antibodies on lung viral load and body weight as a clinical symptoms of disease *in vivo*^46–48^, AvGn-B has emerged as one of a few antibodies with verifiable data demonstrating reduction in disease severity at a histological level *in vivo*. Furthermore, most of these other *in vivo* studies were prophylactic not therapeutic studies. Susceptibility and pathogenesis of SARS-CoV-2 infection in the Syrian hamster is well-characterized^37,38,49^, and this species has been used in SARS-CoV-2 countermeasures studies due to its demonstrated use as suitable experimental model given the close resemblance in pulmonary pathology and clinical features to those observed in human COVID-19 patients^39,47^. The use of this model to assess possible therapeutics is supported by marked reduction of pulmonary monocyte-macrophage infiltration, which may contribute to disease, in the AvGn-B High group. Further, viral load and pulmonary pathology were markedly decreased in Syrian hamsters administered AvGn-B at both doses (Fig. 2).

With extensive pulmonary surface area exposed to SARS-CoV-2 in infected humans and generation of robust amounts of reactive oxygen species (ROS), phospholipid peroxidation occurs, which results in extensive damage to the lungs in experimental H5N1 infection and may contribute to COVID-19 pathology^50–52^. A key pathway leading to lung damage is the oxidative stress causing accumulation of oxidized phospholipids that augment Toll-like receptor 4 (TLR4) expression and signaling by macrophages that in turn upregulates pro-inflammatory cytokine production. Macrophage-rich inflammatory exudates from seriously ill COVID-19 patients usually contain extensive amounts of oxidized lipids^50^. In vulnerable groups of human patients, particularly in those with atherosclerosis, subsets of hyperactive macrophages may play a significant role in heightening the severity of COVID-19^53^. Macrophages in those patients produce a large variety of pro-atherogenic cytokines and chemokines upon stimulation with oxidized low-density lipoprotein (oxLDL). The activation of macrophages is not limited to the affected tissues, but pro-atherogenic stimuli will induce a population of long-lasting inflammatory monocytes in the circulation of those patients producing a “trained innate immune response”^54^. The immunometabolism of monocyte/macrophage populations in atherosclerotic patients revealed that training of histiocytes with oxLDL can stimulate promoters of key inflammatory and tissue-destructive cytokines especially TNF, IL-6, IL-8 and CD36. Notably, CD36 serves as a major scavenger receptor for recognition and internalization of oxLDL^55^. Decreasing the numbers of activated monocytes in the pulmonary circulation and ameliorating the disease severity in the pulmonary parenchyma as shown in the infected hamsters will likely have a therapeutic benefit in the severe form of COVID-19.

Therefore, AvGn-B may offer a promising treatment option for COVID-19 patients predisposed to atherosclerosis due to underlying conditions of hypertension and diabetes mellitus and thus, a more guarded prognosis of COVID-19^56^. The significantly reduced number of infiltrating macrophages in lungs of infected hamsters treated with AvGn-B, support further investigation into its use as both a prophylactic and treatment option for cases of COVID-19 in particularly susceptible patients^57^. Limiting the hyperinflammatory response (*e.g*., cytokine storm) to SARS-CoV-2 infection is another potential therapeutic benefit of AvGn-B treatment. Since severe morbidity and mortality in COVID-19 patients who mount cytokine storm responses involves hyperactivation of the monocyte-macrophage system, AvGn-B may alleviate morbidity and mortality in those patients by reducing the accumulation of activated monocytes/macrophages within SARS-CoV-2 infected lungs.

Another potential therapeutic benefit of AvGn-B related to reduction in lung monocyte infiltration may revolve around the possible role that hyperactivated monocytes might contribute to coagulation and activation of polymorph nuclear leukocytes (PMN) in affected lungs. Microthrombi of the lungs, brain, heart, kidneys and liver plus limbs have been well-documented in COVID-19 patients^58^. By minimizing the accumulation of hyperactivated macrophages in SARS-CoV-2 infected lungs, AvGn-B might confer a therapeutic benefit by limiting induction of intravascular coagulation that primarily takes place within the microcirculation, including in the lungs, that in turn can lead to acute lung injury and sepsis ^59^.

The impact of AvGn-B in protecting against viral infection in both *in vitro* and *in vivo* systems suggests it is a promising option for prophylactic use and therapeutic use in COVID-19 patients. Consequently, in parallel to the *in vivo* studies described above, antibody engineering of clone AvGn-B has been performed and a panel of more potent variants has now been isolated. As shown in Table 2, four of the characterized clones exhibited marked improvement both in blocking the RBD-ACE2 interaction and in viral neutralization activity *in vitro* compared to the parental AvGn-B clone. Clone AvGn-B-G2, in particular, showed an IC_50_ value of 0.17 ng/mL in the CPE assay, nearly 30-fold lower than the IC_50_ value of AvGn-B. In addition, both AvGn-B and its more potent variants, including AvGn-B-G2, recognize the RBD variants, W436R, R408I, N354D, V367F, and N354D + D364Y equally as well as the original RBD from the Wuhan SARS-CoV-2 strain (data not shown) suggesting it should prove effective against escape mutants of SARS-CoV-2.

**Table 2.** Properties of the AvGn-B variants.

| Clone | SARS-CoV-2 RBD<br>(EC<sub>50</sub>, nM) | SARS-CoV-2 RBD<br>(IC<sub>50</sub>, nM) | Neutralizing Activity in CPE<br>(IC<sub>50</sub>, µg/mL) |
| --- | --- | --- | --- |
| AvGn-B | 0.17 | 2.20 | 0.005-0.008 |
| AvGn-B-G2 | 0.09 | 0.26 | 0.00017 |
| AvGn-B-G4 | 0.11 | 0.39 | 0.00064 |
| AvGn-B-H1 | 0.12 | 0.39 | 0.00116 |
| AvGn-B-H2 | 0.09 | 0.39 | 0.00112 |

Future studies will explore the efficacy of the AvGn-B and/or its variants in non-human primate models, several of which have been described for use in SARS-CoV-2 pathogenesis and countermeasure development studies^40,60,61^. Larger studies will also be conducted to examine whether AvGn-B and/or its variants by reducing viral load and accumulation of macrophages within the lung will prevent downstream inflammatory and coagulation sequalae of SARS-CoV-2 infection within other parenchymatous organs, particularly heart, kidneys and liver. Should AvGn-B and/or its variants advance to clinical trials in human patients, its effect on Kawasaki-like disease (K_D_) in SARS-CoV-2 infected children merits clinical investigation. There is accumulating evidence that the monocyte/macrophage system releases cytokines that directly lead to vascular endothelial damage during acute K_D_^62,63^. Investigating the role of AvGn-B and/or its variants in suppressing the cytokine storm by suppressing a pivotal player, monocyte-macrophage system will be important.

Following the identification of SARS-CoV-2 as the causative agent of COVID-19 in early 2020, identification and approval of preventive and therapeutic options for disease management have been a global priority. Using a fully human antibody library, we isolated multiple anti-SARS-CoV-2 spike RBD clones capable of blocking interaction of SARS-CoV-2 spike protein with human ACE-2 receptor and characterized *in vitro* and *in vivo* viral neutralization capacity of clone AvGn-B. Treatment of hamsters with AvGn-B following infection with SARS-CoV-2 resulted in reduced viral load and pulmonary pathology, as evidenced by reduction of macrophage infiltrates. Future studies will confirm whether the increased potency of the AvGn-B-G2 variant in the CPE assay *in vitro* is also shown *in vivo* (*e.g*., lower effective dose). The impact of AvGn-B in protecting against viral infection in both *in vitro* and *in vivo* systems suggests AvGn-B and/or its variants are promising options for prophylactic use and therapeutic use in COVID-19 patients.

## Acknowledgements

The authors thank Lon Kendall for providing Syrian hamsters. This work was supported by the CSU Office of the Vice President for Research (TS) and NIAID R01 AI140442 (TS). AF was supported by the National Institutes of Health (grant 4T32OD010437-18). JL was supported by National Science Foundation (grant 2033260).

